# High variation in handling times confers 35-year stability to predator feeding rates despite altered prey abundances and apparent diet proportions

**DOI:** 10.1101/2022.02.16.480773

**Authors:** Mark Novak

## Abstract

Historical resurveys of ecological communities are important for placing the structure of modern ecosystems in context. Rarely, however, are snapshot surveys alone sufficient for providing direct insight into the rates of ecological processes that underlie how communities function, either now or in the past. In this study, I used a statistically-reasoned observational approach to estimate the feeding rates of a New Zealand intertidal predator, *Haustrum haustorium*, using diet surveys performed at several sites by Robert Paine in 1968–9 and by me in 2004. Comparisons between time periods reveal a remarkable consistency in *H. haustorium*’s prey-specific feeding rates, which contrasts with the changes I observed in prey abundances, *H. haustorium*’s body size distribution, and the proportional contributions of *H. haustorium*’s prey to its apparent diet. Although these results imply accompanying and perhaps adaptive changes in *H. haustorium*’s prey preferences, they are nonetheless anticipated by *H. haustorium*’s high range of variation in prey-specific handling times that dictate not only its maximum possible feeding rates but also the probabilities with which feeding events may be detected during diet surveys. Similarly high variation in detection times (i.e. handling and digestion times) is evident in predator species throughout the animal kingdom. The potential disconnect between a predator’s apparent diet and its actual feeding rates suggests that much of the temporal and biogeographic variation that is perceived in predator diets and food-web structures may be of less functional consequence than currently assumed.

## Introduction

Historical resurveys of ecological communities provide an important means to document com- munity change and contextualize the state of modern ecosystems (Moritz *et al*., 2008; Tingley *et al*., 2009; Chen *et al*., 2009; Sorte *et al*., 2017). Although such resurveys typically involve the comparison of only pairs of points in time, their advantages include the ability to quantify change relative to time periods before the onset of time-series monitoring, which rarely extends prior to the 1970s (Hughes *et al*., 2017; Kuebbing *et al*., 2018). Overall, many historical resurveys have documented substantial changes in community structure (i.e. species composition and abundances); changes that are often, but not always, attributable to climate change, land use, and other, more direct human impacts (Rowe & Terry, 2014; Perry *et al*., 2005; Riddell *et al*., 2021).

Rarely, however, is it possible to use such snapshot surveys to go beyond the characterization of community structure to quantify the rates of the biological processes that underlie how communities function, such as growth, predation, and competition (Paine, 1966; McCoy & Pfister, 2014; Urban *et al*., 2016). Studies in which this has been possible have revealed sometimes unexpected insights. For example, Rowe *et al*. (2011) combined historic and modern surveys of small mammal species and their body-size distributions with metabolic scaling laws to relate changes in community structure to marked declines in rates of total energy use within Great Basin communities since the late 1920s. These patterns, however, contrasted markedly with the findings of Terry & Rowe (2015) who used the same approach to reveal that, despite substantial changes in small mammal body-size distributions and community structure, total energy use remained stable over the period of rapid climate warming that occurred at the terminal Pleistocene. Studies that quantify process rates can therefore provide levels of insight into underlying drivers of change or stasis that surveys of community structure alone may miss.

Unfortunately, most survey studies that quantify process rates have had to rely on species-agnostic theory or empirical relationships (such as metabolic scaling laws) or have depended on the existence of parameter-rich physiology-based models. For example, Atcheson *et al*. (2012) used a bioenergetic model to combine estimates of apparent diet and prey availability with estimates of individual growth rates from scale circuli to simulate and compare rates of prey biomass consumption by Steelhead fishes over 18 years in the North Pacific. Although the mechanistic basis and structural assumptions of such models are often well-grounded and empirically-validated by applications in present-day settings, their appropriateness to historical time periods can be difficult to affirm or rely upon given compounding estimation uncertainties and the pace by which evolutionary and other biological changes (e.g., behavioural plasticity) can proceed.

In this study, I used an alternative, statistically-reasoned approach to directly estimate and assess changes in the prey-specific feeding rates of a predatory intertidal whelk, *Haustrum haustorium*, whose diet was surveyed at several northern New Zealand sites by Robert (Bob) T. Paine (Estes *et al*., 2016; Dayton *et al*., 2016; Palumbi *et al*., 2017; Power *et al*., 2018) in 1968– 1969 and which I resurveyed in 2004. The approach I used to estimate feeding rates contrasts with the aforementioned theory and model-based approaches in minimally requiring data on only two aspects of predator foraging: estimates of a predator’s apparent diet from feeding surveys (i.e. diet proportions) and estimates of feeding events’ detection times (defined below). Based on the 35-year time-span and an expectation of non-equilibrial, dynamically-changing species interactions and abundances in the region (e.g., Benincà *et al*., 2015) and intertidal systems in general (e.g., Katz, 1985; Menge *et al*., 2022; Sorte *et al*., 2017), I naively expected to see a weak correspondence between the feeding rates of the two time periods. Instead, my comparisons revealed a remarkable stability in *H. haustorium*’s prey-specific feeding rates that contrasted with the changes I observed in prey abundances, *H. haustorium*’s body-size distribution, and the proportional contributions of *H. haustorium*’s prey species to its apparent diet. Additional analyses implicated similarly-large changes in *H. haustorium*’s prey-specific per capita attack rates (i.e. its prey preferences).

I recognized the inevitability of *H. haustorium*’s feeding-rate consistency only in hindsight. The underlying mechanism — attributable to the wide range of *H. haustorium*’s handling times across its many prey species — has nonetheless been recognized for over 120 years as the effect of correlated denominators on the correlation of ratios (Pearson, 1897). The results of my analyses thereby speak to the importance of statistical thinking when interpreting survey data, and to the importance of studying ecological process rates rather than community structure alone. They also emphasize the importance of distinguishing between a predator’s apparent and true diet when making temporal (or geographic) comparisons to understand how food webs and communities function.

## Methods

### Data collection

#### Study system

*Haustrum haustorium* is a muricid whelk that is endemic to the North and South Islands of New Zealand (Tan, 2003). Its fossil record shows *H. haustorium* to have grown to 80 mm shell length (Tan, 2003), but in modern times its size rarely exceeds 55 mm (Novak, 2008).^1^ Its diet varies through ontogeny, but primarily consists of herbivorous limpets, chitons and snails, filter-feeding barnacles and mussels, and its congener *H. scobina* (formerly *Lepsiella scobina*) with whom it shares many prey species (Luckens, 1975; McKoy, 1969; Morton & Miller, 1968; Ottaway, 1977; Patrick, 2001; Walsby, 1977; Novak, 2008; 2010; 2013). *H. haustorium* drills through the shells of its prey and/or flips them over to digest and ingest the “soup” through its extended proboscis (Fig. 1). A feeding event can last hours to more than a day and thus through one or more low-tide periods depending on the temperature, the prey’s identity, and the sizes of the whelk and prey individual (Novak, 2010; 2013).

**Figure 1:**
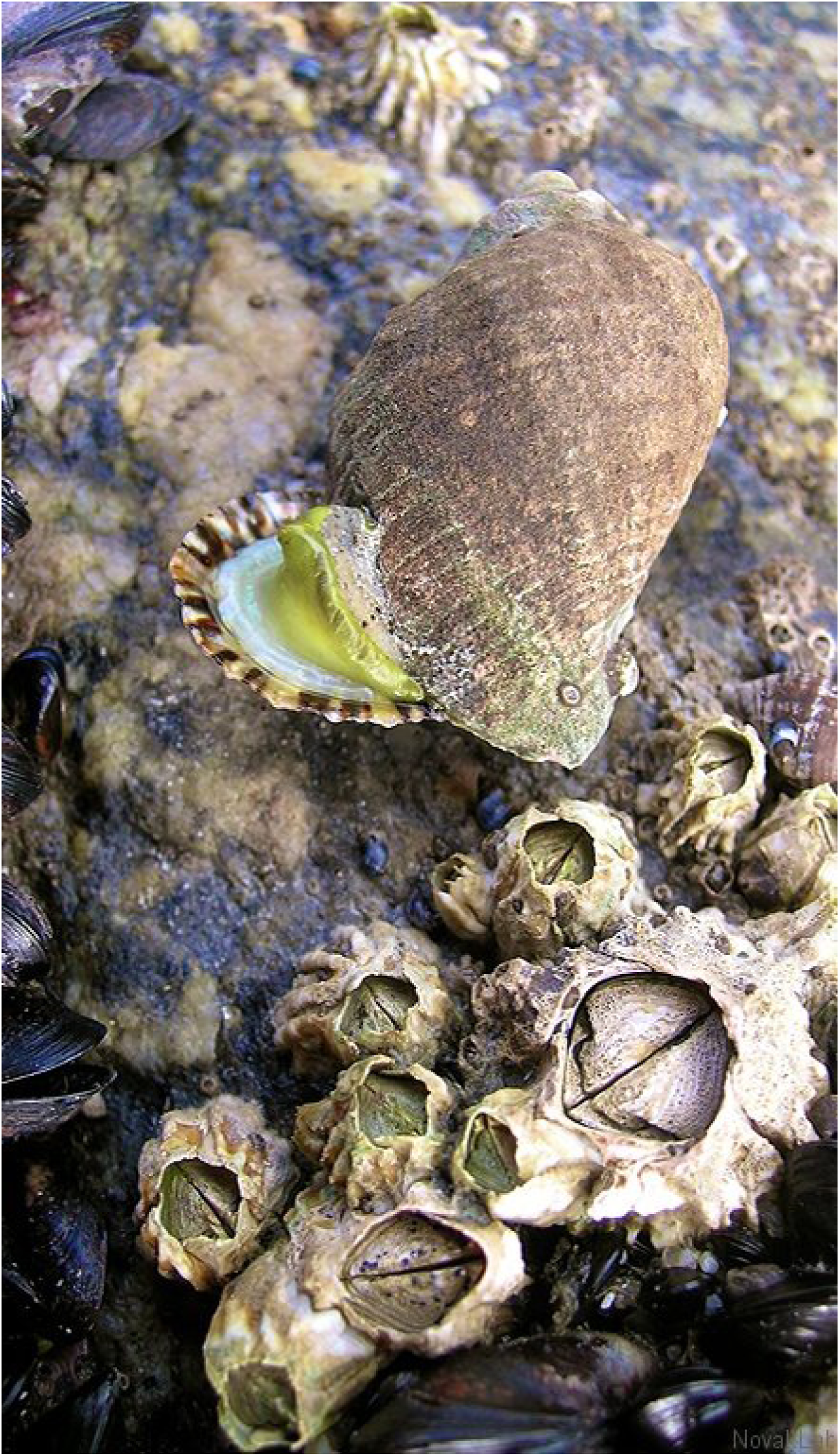
*Haustrum haustorium* feeding on the limpet *Cellana ornata*, surrounded by additional prey species: *Xenostrobus pulex* mussels, *Epopella plicata* and *Chamaesipho columna* barnacles, *Austrolittorina antipodum* snails, and its congeneric intraguild prey, *H. scobina* (center right).

#### Feeding surveys

Feeding surveys during low-tide periods are a standard means to determine the apparent diet of whelks and many other intertidal predators (e.g., Paine, 1963; Menge, 1974; Hughes & Burrows, 1991; Yamamoto, 2004). They consist of a systematic search of an area of rocky shore, carefully inspecting each found individual to determine whether or not it is feeding, measuring its shell length (*±* 1 mm) and, if it is feeding, identifying and measuring the size of its prey. Paine conducted such surveys at ten sites along the northern coast of the North Island between November 1968 and May 1969 (Table S1). In June 2004, using Paine’s site names, descriptions, and hand-drawn maps, I was able to relocate and access five of the same sites to resurvey *H. haustorium*’s diet using the same protocols.

**Table 1:** Summary of Paine’s 1968-9 and my 2004 feeding observations. Observations refers to the total number of whelks inspected. % feeding refers to the proportion of observed whelks that were feeding. Parentheticals are the binomial confidence interval (95% coverage probability) calculated using the Wilson method.

| Site | Observations |  | % feeding |  |
| --- | --- | --- | --- | --- |
|  | 1968-9 | 2004 | 1968-9 | 2004 |
| Leigh - Echinoderm Reef | 228 | 72 | 13.6 (9.7-18.7) | 13.9 (7.7-23.7) |
| Leigh - Tabletop Rocks and Boulders | 44 | 268 | 65.9 (51.1-78.1) | 5.2 (3.1-8.6) |
| Leigh - Waterfall Rocks | 275 | 1060 | 17.5 (13.4-22.4) | 10.1 (8.4-12.1) |
| Rangitoto Island - Whites Beach | 93 | 130 | 11.8 (6.7-19.9) | 20.8 (14.7-28.5) |
| Red Beach - Whangaparaoa | 461 | 37 | 24.5 (20.8-28.6) | 5.4 (1.5-17.7) |
| Sum/Average | 1101 | 1567 | 26.7 | 11.1 |

#### Prey abundance surveys

Paine also conducted abundance surveys of *H. haustorium*’s prey species at several sites, including three of the sites where he performed feeding surveys and which I was able to resurvey (Table S1). Abundance surveys entailed the use of a 0.3*×*0.3 m quadrat which Paine placed randomly at 15 positions along a transect line (of unknown length) located haphazardly within the same area in which feeding surveys were subsequently conducted. All mobile prey species within the quadrats were counted. I repeated these surveys using 15 quadrats positioned randomly along a 20 m transect. Paine often distinguished among tidal zones (e.g., the “oyster zone” and “1 ft. above *Xiphophora* zone”), surveying a transect (or two) in each of them. I matched my survey areas to these zones as best I could, though sometimes zonation patterns were not as clear as they had apparently been for Paine.

#### Species identifications

Three things are worth noting in regards to species identifications of key taxa:

(i) The whelk referred to as *Neothais scalaris* in the only paper that Paine published of his New Zealand work (Paine, 1971) is now called *Dicathais orbita*. Among its differences from *H. haustorium* is that *Dicathais* occurs on more exposed shores where its apparent diet consists primarily of *Perna* mussels.
(ii) Although Paine (1971) mentions having observed *Dicathais* at multiple (unspecified) sites, and to have estimated its density to be 17 m*^−^*^2^ at Red Beach, Whangaparoa Peninsula, specifically, I observed few to no *Dicathais* at the sites which I resurveyed, including the Red Beach, Whangaparoa Peninsula site that I surmised Paine to have surveyed for *H. haustorium*. I nonetheless consider it unlikely that Paine mistook small *H. haustorium* or *Paratrophon* spp. — which can appear similar to small *Dicathais* and which I did observe at Red Beach — for *Dicathais*.
(iii) It is possible that the prey species *H. scobina* reported on here (and in Novak (2010; 2013) for sites around the South Island) is conflated with the sister taxon *H. albomarginatum* (Barco *et al*. 2015; but see Tan 2003; O’Mahoney 2020).

### Data analysis

#### Estimating feeding rates

The approach I used for estimating *H. haustorium*’s prey-specific feeding rates from diet surveys appears to have been first used by Charles Birkeland (Birkeland, 1974) who obtained his Ph.D. with Paine as primary advisor. It was re-derived by Novak *et al*. (2017) and ostensibly several others (Bajkov, 1935; Englund & Leonardsson, 2008; Speirs *et al*., 2000; Woodward *et al*., 2005). The approach relies on the following information:

(i) the count of the number of predator individuals that, in the course of a snapshot diet survey, are observed to be feeding on each focal prey species (*n_i_*);

### the count of the total number of predator individuals that are surveyed (*n*); and

(i) an estimate of the (average) length of time over which a feeding event on each focal prey species remains detectable to an observer (*d_i_*).

A formal derivation is summarized as follows: Consider a generalist predator population whose diet consists of *i* = 1*, . . . , S* different prey species on which predator individuals feed only one prey item at a time. If *f_i_* is the predator population’s average feeding rate on the *i^th^* prey species (which we wish to estimate) then, over some time period *T* , an average individual will consume *f_i_T* individuals of prey *i*. If each of these feeding events remains detectable to an observer for time *d_i_* then the total time that the predator individual could have been seen feeding in time period *T* is *f_i_d_i_T* and the proportion of time it could have been seen feeding on prey *i* is *f_i_d_i_*. It follows that, if we perform a snapshot feeding survey of *n* independent and equivalent predator individuals, the expected proportion of individuals we should observe feeding on each prey species, *p_i_*, will also be *f_i_d_i_*. Therefore, and since the maximum likelihood estimator of *p_i_* is *n_i_/n*, we can estimate prey-specific feeding rates as

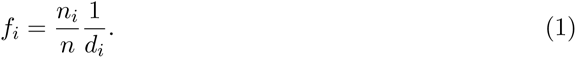

In using the approach we make no assumptions regarding the form of the predator’s functional response and need not know prey nor predator abundances.

Clearly, the primary challenge for applying the approach to diet surveys is to have information on detection times. Indeed, depending on its detection time, a species that is frequently observed in a predator’s apparent diet may in fact be only infrequently consumed by the predator if its detection time is long (Novak, 2010; Fairweather & Underwood, 1983). I estimated *H. haustorium*’s prey-specific detection times (in days) on the basis of extensive laboratory experiments which I had previously performed for *H. haustorium* populations of New Zealand’s South Island (Novak, 2013). These experiments involved placing individuals of varied sizes into isolated aquaria, providing them focal prey of varied sizes and identities, and subsequently classifying each whelk as either feeding or not feeding on a near hourly basis or continuously with video surveillance. Whelk and prey size combinations maximized or exceeded the range of relative sizes observed in the field. The temperature was varied between 10 and 18 *^◦^*C by placing the aquaria in temperature-controlled rooms. For each prey species, I regressed the observed detection times on whelk size, prey size, and temperature (all variables log*_e_*-transformed, see Novak, 2013) and used the resulting regression coefficients to back-calculate the expected detection time of each feeding event that Paine and I had observed in the field. In doing so, I used the mean water temperature measured in the given year and month at the Leigh Marine Laboratory for all surveyed sites (Evans & Atkins, 2013; Costello, 2015), the laboratory being centrally located to all sites and providing the only *in situ* temperature record that extends to the 1960s. Prey for which I had not estimated detection-time regression coefficients in the experiments were matched to the most similar species for which they had been estimated (Table S2). Feeding observations in which either the size of the prey or whelk were unknown (typically because the prey was “swallowed” when the predator closed its operculum too quickly) were assigned the species’ mean detection time across all observations.

#### Comparisons of 1968–9 & 2004

I used several measures of correlation and deviation to quantify the similarity of feeding rates between 1968–9 and 2004. I ignored prey species that were not observed in *H. haustorium*’s diet at a given site in both time periods and used all remaining time-period pairs of site-specific prey species from across all five sites to calculate similarities (see *Supplementary Materials* for analyses including species observed in only one time period). As is typically the case (e.g., Preston *et al*., 2019), feeding rates varied over several orders-of-magnitude and exhibited a right-skewed frequency distribution due to their underlying multiplicative nature. I therefore calculated the correlation between time periods in three ways: using Pearson’s linear correlation coefficient on the natural scale (*r*), using Pearson’s correlation coefficient after log_10_-transformation (*r*_10_), and using Spearman’s rank correlation coefficient (*r_s_*). I estimated *p*-values using two-sided tests. I also calculated the mean logarithmic difference (*MLD*) and the mean absolute logarithmic difference (*MALD*) between feeding-rate pairs, these both being measures of relative similarity (since log_10_(*x*) *−* log_10_(*y*) = log_10_(*x/y*)). I repeated these same calculations for the prey-specific diet proportions (*p_i_* = *n_i_/n*) and the field-calculated detection times (*d_i_*), restricting these comparisons to the same site-prey pairs that were included in the comparison of the feeding rates.

In order to determine whether (dis)similarities between time periods in any of the just-mentioned three variables were associated with changes in *H. haustorium*’s or its prey’s sizes, I plotted histograms of whelk and prey sizes and formally assessed differences between time periods using two-sided Kolmogorov-Smirnov (KS) tests. I also used multiple linear regression to regress whelk size (log*_e_*-transformed) on prey size (log*_e_*-transformed), time period, and their first-order interaction to determine whether there was a change in *H. haustorium*’s prey-size selectivity.

Finally, in order to determine whether (dis)similarities between time periods in *H. haustorium*’s feeding rates were associated with changes in its prey preferences, I used the estimator derived by Novak & Wootton (2008) and clarified by Wolf *et al*. (2017) to calculate *H. haustorium*’s per capita attack rates. This estimator uses the same information as used to estimate feeding rates (i.e. the *n_i_* prey observations and *d_i_* detection times), but also makes use of the number of surveyed individuals that are observed to be *not* feeding (*n*_0_), requires knowledge of each prey’s abundance (*N_i_*), and necessitates the specification of a functional-response model (Novak *et al*., 2017). I assumed the multi-species extension of the Holling Type II functional response (e.g., Murdoch, 1973) and that *H. haustorium*’s handling times equaled its detection times (i.e. *h_i_*= *d_i_*). Under these assumptions, which are well-justified for *H. haustorium* (see Novak, 2010; 2013; Novak *et al*., 2017), the estimator for *H. haustorium*’s per capita attack rate on prey *i* is

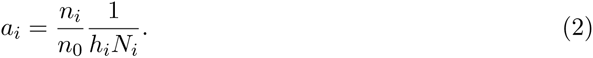

In absolute terms, these per capita attack rates represent the number of prey eaten per predator per day per available prey, with abundances represented by densities (here per square meter). In relative terms, they represent the predator’s prey preferences accounting for differences in prey handling times and abundances (Novak & Wootton, 2008).

Because the attack rate estimator requires estimates of prey abundances, I calculated attack rates only for the subset of three sites where both Paine and I had estimated these using quadrat surveys. I then calculated the between time-period correlations and deviations of the attack rates, feeding rates, diet proportions, detection times, and prey abundances for these sites just as described above. Finally, I used multiple linear regression to regress feeding rates (log*_e_*-transformed) on prey abundances (log*_e_*-transformed), time period, and their first-order interaction to determine whether there was an effect of time period on the density dependence of *H. haustorium*’s feeding rates (i.e. its across-species “functional response”).

## Results

### Feeding survey sites

Across the five sites at which both Paine and I performed feeding surveys, Paine observed 232 of 1101 total individuals feeding on 10 different species (Table 1). In my resurveys, I observed 160 of 1567 total individuals feeding on 16 different species. Across sites, the proportions of feeding individuals ranged from 11.8 to 65.9% for Paine and from 5.2 to 27.3% for me. Paine observed *H. haustorium* feeding on 2 species that I did not observe whereas I observed it feeding on 8 species that Paine did not, therefore together we observed *H. haustorium* feeding on 18 different prey species.

There were 7 species on which both Paine and I observed *H. haustorium* feeding at the same site. For these 7 species, there were 17 site-species feeding-rate pairs for me to compare between 1968–9 and 2004 (Fig. 2A). These varied over two orders of magnitude (from 1.71 *·* 10*^−^*^3^ to 0.51 *·* 10*^−^*^1^ prey per predator per day), were positively correlated between time periods for all three correlation measures (*r* = 0.58*, p* = 0.01; *r*_10_ = 0.79*, p <* 0.001; *r_s_*= 0.78*, p <* 0.001), and tended to be greater in 1968–9 than in 2004 (mean deviation and 95% bootstrapped confidence interval: *MLD* = 0.220 (0.020, 0.418), *MALD* = 0.404 (0.294, 0.520)).

**Figure 2:**
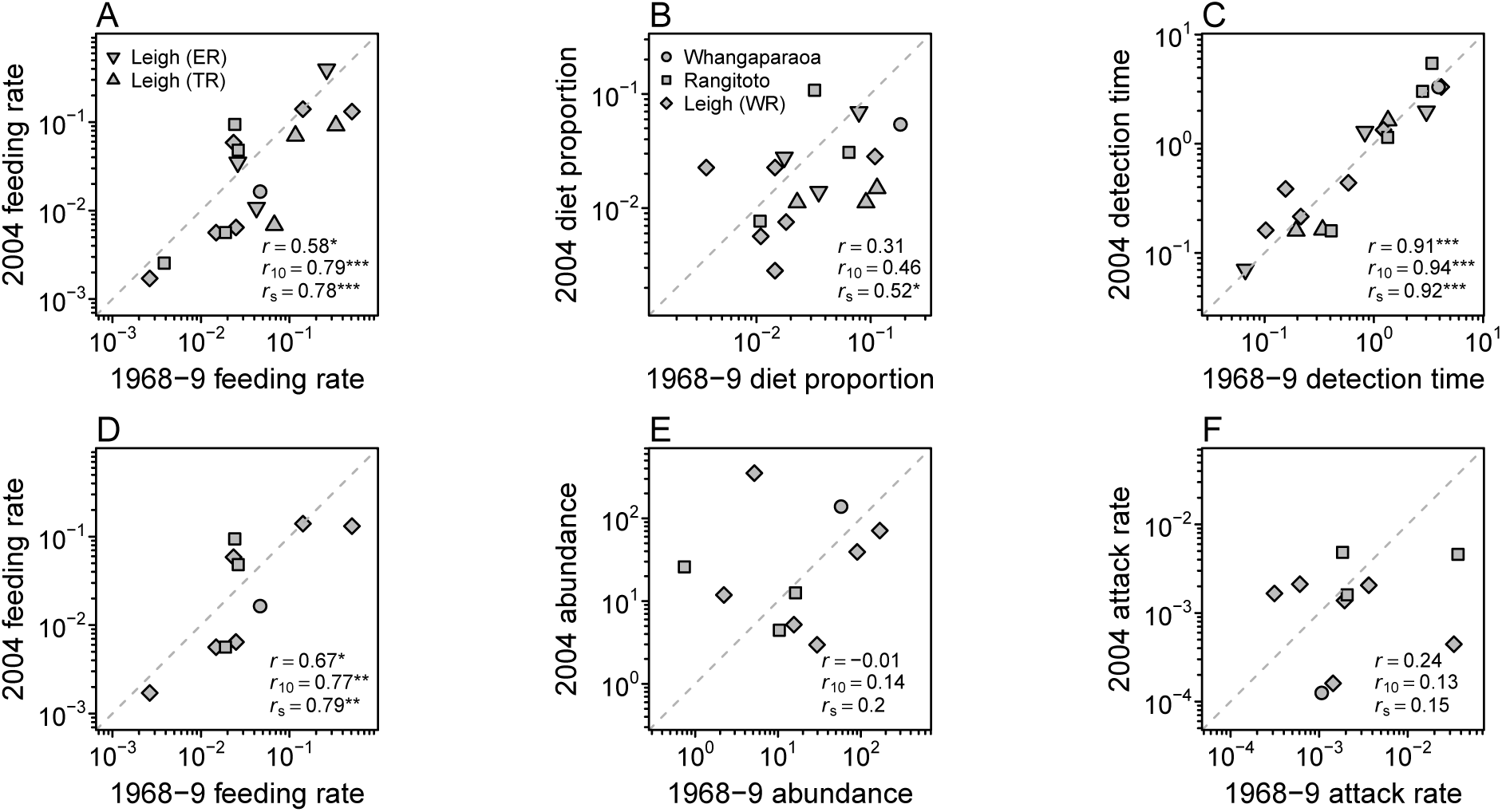
The between time-period correlation of prey-specific (A) feeding rates, (B) apparent diet proportions, and (C) detection times among all sites where Paine and I surveyed *H. haustorium*’s diet, and of prey-specific (D) feeding rates, (E) abundances, and (F) per capita attack rates for the subset of sites where Paine and I also surveyed prey abundances. I calculated three correlations for each comparison to assess the linearity and monotonicity of the time-period (dis)similarities: Pearson’s correlation (*r*), Pearson’s correlation after log_10_-transformation (*r*_10_, as plotted), and Spearman’s rank correlation (*r_s_*). The probability of observing a correlation at least as extreme as the observed correlation under the null hypothesis of no correlation (two- tailed test) is indicated by asterisks: *** *p <* 0.001; ** *p <* 0.01; * *p <* 0.05; otherwise *p >* 0.1.

In contrast to the feeding rate, *H. haustorium*’s apparent diet proportions showed relatively little similarity between time periods (Fig. 2B). That is, although the diet proportions exhibited similar variation within each time period (varying from 2.83 *·* 10*^−^*^3^ to 1.8 *·* 10*^−^*^1^), their between time-period correlations were lower and less clearly different from zero (*r* = 0.31*, p* = 0.22; *r*_10_ = 0.46*, p* = 0.06; *r_s_* = 0.52*, p* = 0.03). They also tended to be greater in 1968–9 than in 2004 (*MLD* = 0.233 (0.018, 0.439), *MALD* = 0.433 (0.313, 0.599)). On the other hand, mean detection times were very similar between time periods (Fig. 2C). These varied over two orders of magnitude (from 1.6 to 130.8 hours), were highly correlated between time periods for all three measures (*r* = 0.92*, p <* 0.001; *r*_10_ = 0.94*, p <* 0.001; *r_s_* = 0.92*, p <* 0.001), and were not distinguishable between time periods (*MLD* = 0.011 (*−*0.079, 0.100), *MALD* = 0.149 (0.094, 0.210)). Although *H. haustorium*’s size range was unchanged between time periods, its size distribution showed a clear shift towards smaller individuals in 2004 relative to 1968–9 (Fig. 3; 1968-9: 9.8 *−* 63.0 mm, *x̄* = 34.7 mm; 2004: 9.0 *−* 62.0 mm, *x̄* = 30.0 mm; KS test: *D* = 0.30, *p <* 0.001, all five sites combined). The size distribution of prey individuals was also shifted towards smaller individuals in 2004 (Fig. 3; 1968-9: 1.0*−*36.0 mm, *x̄* = 16.5 mm; 2004: 2.0*−*28.0 mm, *x̄* = 10.2 mm; KS test: *D* = 0.55, *p <* 0.001). *H. haustorium*’s relative prey-size selectiv- ity, however, appeared unchanged between time periods, with multiple regressions providing no support for main or interactive effects of time period (Fig. 3, Tables S3-S5, log*_e_*(*Predator size*) = 2.34+0.46*·* log*_e_*(*Prey size*), *F*_1_,_381_ = 629.9, *p <* 0.001, *R*^2^ = 0.62 for both periods combined).

**Figure 3:**
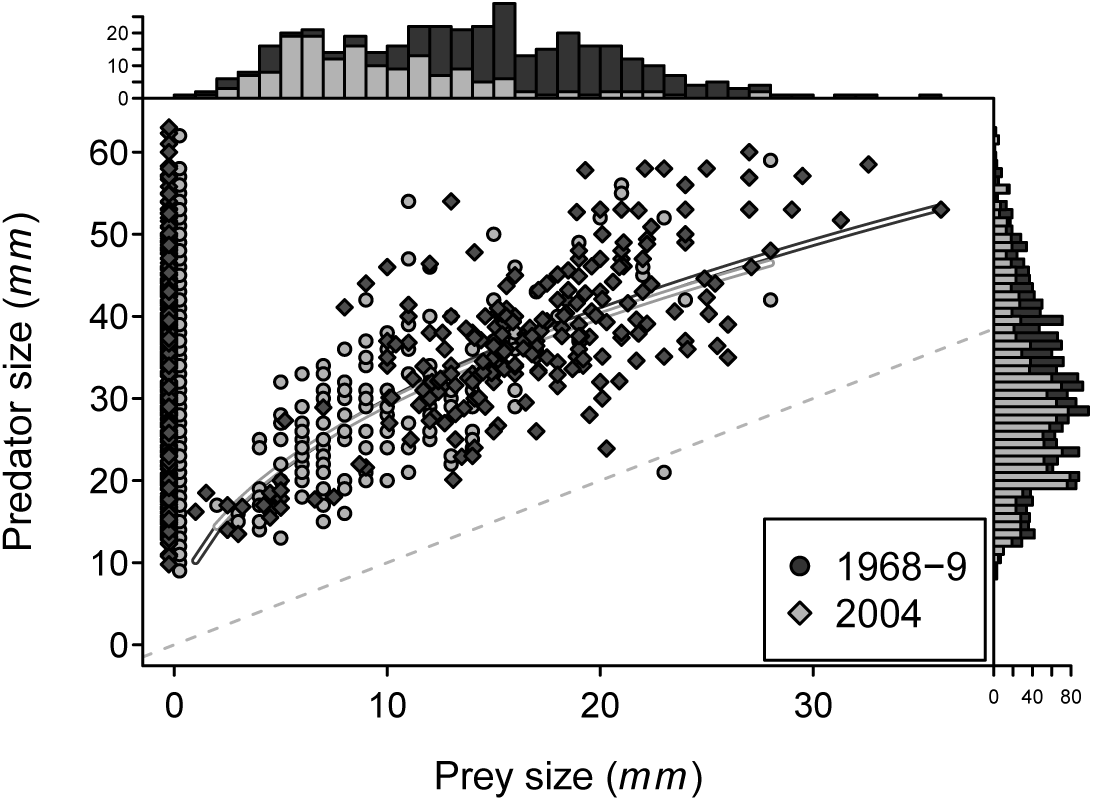
Although the sizes of *H. haustorium* individuals and the sizes of their prey individuals were smaller in 2004 than in 1968–9, *H. haustorium*’s size-selectivity was unchanged between time periods. See Tables S3-S5 for regression summaries. The values near a prey size of 0 mm indicate the sizes of non-feeding whelks and are omitted from the prey-size frequency histogram. Note that this figure includes the *H. haustorium* and prey individuals of all observations made at the five focal study sites (rather than just the subset of temporally-paired prey-specific estimates considered in Fig. 2).

### Feeding and abundance survey sites

Feeding rates were even more clearly similar between time periods for the 10 pairs of site-species estimates (6 prey species) from the three sites where Paine and I performed both feeding and abundance surveys (Fig. 2D; *r* = 0.67*, p* = 0.03; *r*_10_ = 0.77*, p <* 0.01; *r_s_* = 0.79*, p <* 0.01; *MLD* = 0.15 (*−*0.116, 0.394), *MALD* = 0.40 (0.283, 0.508)). Just as seen when considering all five sites, the between time-period similarity of the apparent diet proportions was lower (not shown; *r* = 0.35*, p* = 0.32; *r*_10_ = 0.51*, p* = 0.13; *r_s_* = 0.62*, p* = 0.053; *MLD* = 0.147 (*−*0.165, 0.416), *MALD* = 0.447 (0.322, 0.574)), but the similarity of mean detection times was high (not shown; *r* = 0.90*, p <* 0.001; *r*_10_ = 0.93*, p <* 0.001; *r_s_* = 0.87*, p* = 0.003; *MLD* = *−*0.006 (*−*0.138, 0.123), *MALD* = 0.161 (0.085, 0.250)).

Prey abundances varied over two orders of magnitude within both time periods (varying from 0.74 to 351 individuals per m^2^), but showed no correspondence between time periods (Fig. 2E; *r* = *−*0.009*, p* = 0.98; *r*_10_ = 0.14*, p* = 0.71; *r_s_* = 0.2*, p* = 0.58; *MLD* = *−*0.178 (*−*0.746, 0.338), *MALD* = 0.718 (0.420, 1.073)). This was similarly true for the estimates of *H. haustorium*’s per capita attack rates, which also varied over three orders of magnitude within time periods (varying from 5.2 *·* 10*^−^*^6^ to 1.5 *·* 10*^−^*^3^ prey per predator per day per prey available) but showed no correspondence between time periods (Fig. 2F; *r* = 0.24*, p* = 0.50; *r*_10_ = 0.13*, p* = 0.72; *r_s_* = 0.15*, p* = 0.68; *MLD* = 0.348 (*−*0.116, 0.835), *MALD* = 0.685 (0.402, 1.017)).

Regressing feeding rates on prey abundances did not show main or interactive effects of time period on the density dependence of *H. haustorium*’s across-species “functional response” (Tables S6-S7), with the simpler model combining time periods revealing that feeding rates increased with a decelerating rate as prey abundances increased (Fig. 4, Table S8, log_10_ *f_i_* = *−*2.26 + 0.52 *·* log_10_ *N_i_*, *F*_1_,_23_ = 8.41, *p* = 0.008, *R_adj_*^2^ = 0.24).

**Figure 4:**
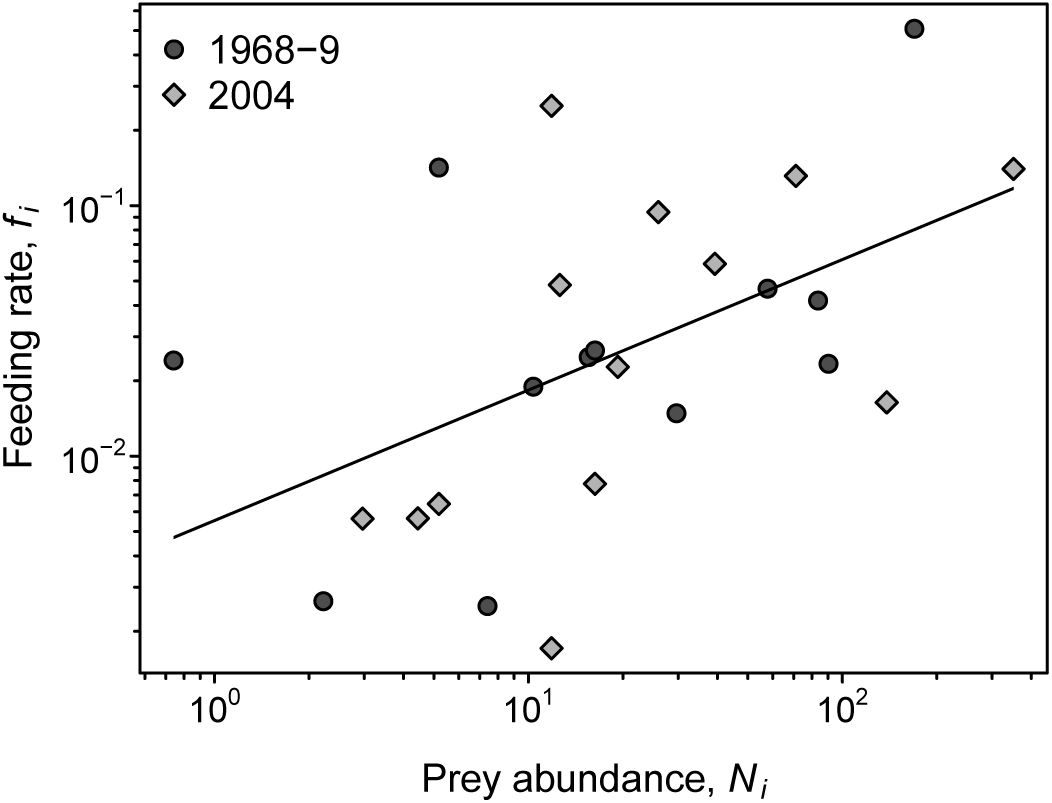
*H. haustorium*’s prey-specific feeding rates (prey eaten per predator per day) increased as a decelerating function (logarithmic slope *<* 1) of prey abundance (per m^2^) and were not distinguishable by time period (Tables S6-S8). Note that this regression includes five temporally- unpaired estimates that reflect feeding rate and abundance estimates for prey species which only Paine *or* I observed (rather than just the subset of temporally-paired prey-specific estimates considered in Fig. 2).

## Discussion

That feeding rates are dynamic and respond to many aspects of a predator’s environment is a central, well-supported thesis. The importance of predator and prey abundances, their body sizes, and environmental temperature has elicited particularly strong research attention within the vast literatures relating to predator foraging ecology, food webs, and the impacts of climate change. Although water temperatures in northern New Zealand have not exhibited a systematic trend to date (Shears & Bowen, 2017), my resurveys of Bob Paine’s study sites revealed little similarity in *H. haustorium*’s apparent diet between 1968–9 and 2004. My resurveys further showed an overall reduction in *H. haustorium*’s body size which, though not associated with changes in the *relative* size of chosen prey individuals, was accompanied by substantial changes in community structure. These changes in apparent diet proportions and prey abundances inferred by my main analyses are corroborated by additional comparisons that included (rather than excluded) species observed by only Paine or only me (see *Supplementary Materials*).

Given these observations and their consistency with the dynamic nature of rocky intertidal systems in the region (e.g., Benincà *et al*., 2015) and the world more generally (e.g., Katz, 1985; Menge *et al*., 2022; Sorte *et al*., 2017), I expected *H. haustorium*’s prey-specific feeding rates to have been similarly altered in the 35 years that separated Paine’s and my surveys. Instead, as estimated by a statistically-reasoned approach that does not rely on species-agnostic scaling laws, parameter-rich energetic models, or even the specification of a particular functional-response model, *H. haustorium*’s feeding rates showed a remarkable stability between the two time periods (Fig. 2A,D). That is, although feeding rates were overall higher in 1968–9 than in 2004 (possibly due to the change in *H. haustorium*’s body size), prey-specific feeding rates evidenced a high degree of temporal consistency in their relative within time-period magnitudes regardless of the metric of similarity I employed.

On the face of it, this contrast between *H. haustorium*’s feeding-rate stability versus the changes in its prey’s abundances and apparent diet contributions implies a substantial compensatory response in *H. haustorium*’s prey preferences. This inference was underscored by my comparison of *H. haustorium*’s per capita attack rates at the subset of sites where these could be estimated assuming a multi-species Type II functional response (for which the attack-rate parameters encapsulate prey preferences). That is, regardless of how their similarity was quantified, attack-rate estimates in 1968–9 showed no similarity to the estimates of 2004 (Fig. 2F). Indeed, the temporal consistency of the saturating (albeit loose) relationship between *H. haustorium*’s feeding rates and its prey’s abundances (Fig. 4) that was associated with these changes in attack rates could be inferred to indicate an adaptive response in prey preferences to altered prey abundances (*sensu* Abrams, 1999; Kondoh, 2003).

I believe this final inference to be incorrect however. Instead, I attribute the stability of *H. haustorium*’s feeding rates to a mechanism that is statistical in nature and was recognized in 1897 soon after the formal definition of Pearson’s measure of correlation itself.

### The inevitability of feeding-rate stability

Pearson’s correlation coefficient *r* is a measure of the linear association between two variables (Pearson, 1895; Bravais, 1844). Pearson (1897) was the first to note that two ratios (*x/w* and *y/z*) will be correlated when their denominator variables are correlated, even if the numerator variables are entirely uncorrelated. He derived the following expression with which to approximate this expected correlation of ratios using the correlations between each pair of variables and each variable’s coefficient of variation (*v*, its standard deviation divided by its mean):

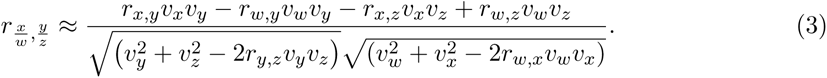

Although it assumes that the coefficients of variation are small (Kim, 1999), and although an exact expectation may be obtained with a permutation-based approach (see *Supplementary Materials*), Pearson’s approximation provides useful insight into how a correlation between ratios will arise. In fact, in the context of understanding the stability of *H. haustorium*’s feeding rates

(i.e. where 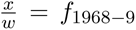 and 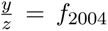), the approximation may be further simplified by (*i*) letting the numerator variables (the *x, y* apparent diet proportions; *n_i_/n* in eqn. 1) and the denominator variables (the *w, z* detection times; *d_i_*in eqn. 1) be uncorrelated with each other within and across time periods (i.e. *r_y,z_* = *r_w,x_*= *r_w,y_* = *r_x,z_* = 0) and (*ii*) letting the coefficients of variation of the two numerator variables and the two denominator variables each be the same across time periods (i.e. *v_n_* := *v_y_*= *v_x_* for the diet proportions and *v_d_*:= *v_z_*= *v_w_* for the detection times). Under these simplifications, Pearson’s approximation is reduced to

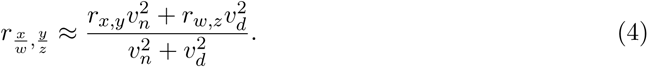

Since the denominator of eqn. 4 simply scales the response between *−*1 and +1, it follows that feeding rates will tend to be positively correlated between time periods whenever the detection times are positively correlated and exhibit a sufficiently large coefficient of variation across prey species, even if the apparent diet proportions are uncorrelated or negatively correlated (Fig. 5). Feeding-rate stability can therefore occur despite substantial changes in the predator’s prey preferences or its prey’s abundances. The same logic applies using Spearman’s rank correlation coefficient since it is just the Pearson correlation of rank-ordered values.

**Figure 5:**
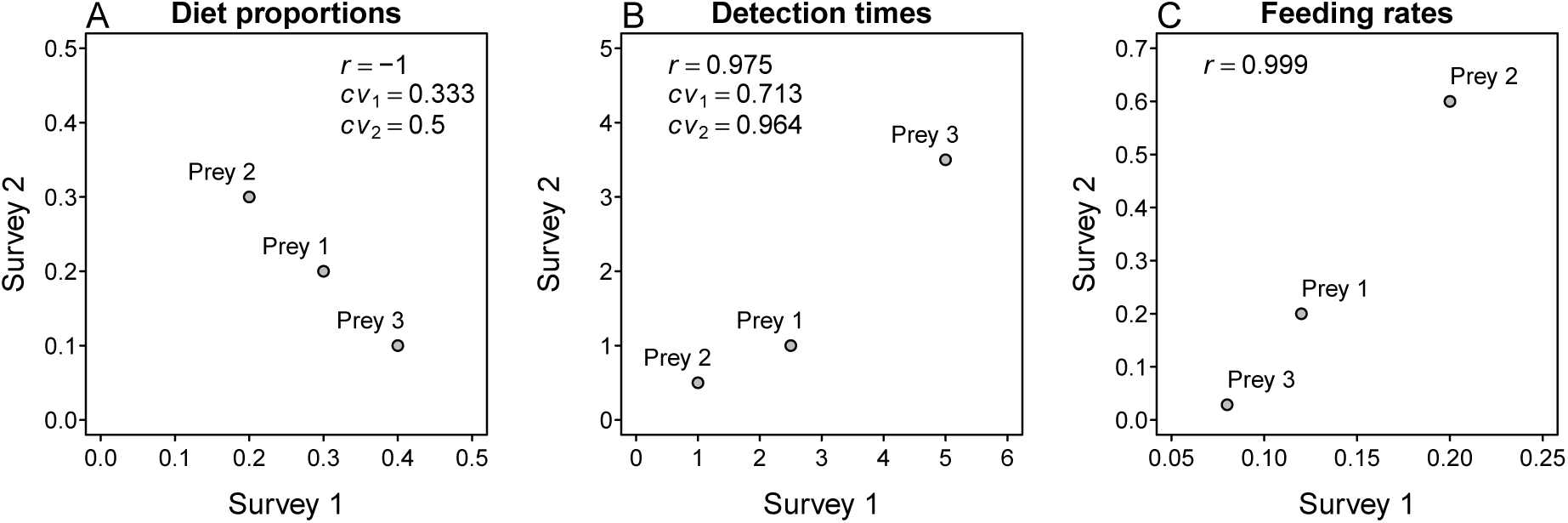
A hypothetical example of the statistical mechanism causing correlated ratios of which Pearson (1897) warned. The panels show two surveys between which a predator’s (A) apparent diet proportions on three prey species are perfectly negatively correlated (*r* = *−*1.00), but its (B) detection times are positively correlated (*r* = 0.975) and exhibit sufficiently high coefficients of variation (*cv*) for its (C) feeding rates to be strongly positively correlated (*r* = 0.999). (Given correlations are exact, not estimated using eqns. 3 or 4.)

Pearson (1897) referred to the non-zero correlation of ratios involving uncorrelated numerator and correlated denominator variables as being spurious (but see Haig, 2003; for discussion of the term itself). When inference is being made regarding the relationship of the two numerator variables the issue is indeed a major problem that has plagued — and continues to plague — diverse scientific disciplines (e.g., Jackson & Somers, 1991; Kenney, 1982; Atkinson *et al*., 2004; Håkanson & Stenström-Khalili, 2009; Williams *et al*., 2021), leading many to infer a relationship between measured variables when in fact none exists. However, as first noted by Yule (1910), the relationship is not spurious when inference is being made regarding the ratios (Aldrich, 1995), as is the case in using eqn. 1 to estimate feeding rates. That is, the correlation of ratios due to correlated denominator variables reflects (the linear aspect of) the true relationship between the ratios themselves. The stability of *H. haustorium*’s feeding rates between the two time periods is therefore not a spurious inference. Instead, it is the inevitable consequence of *H. haustorium*’s positively-correlated and wide-ranging detection times that are themselves a direct consequence of the wide-ranging handling times that *H. haustorium* exhibits across its diverse diet.

### Generality and assumptions

At the species level, *H. haustorium*’s detection times were estimated to vary between 1.6 and 130.8 hours. A wide range of detection times is typical for whelks (e.g., Yamamoto, 2004) and many other taxonomically-diverse consumers — from fishes to birds, seastars, spiders, and flies (e.g., Preston *et al*., 2017; Hilton *et al*., 1998; Uiterwaal & DeLong, 2020; Campos & Lounibos, 2000; Menge, 1972) — and is the consequence of a wide variety of both general and specific prey attributes. These include differences in digestible tissue mass (e.g., acorn barnacles are smaller than mussels), chemical defenses (e.g., *H. scobina* exudes a dark purple substance when consumed by *H. haustorium* (*pers. obs.*) and takes much longer to consume than similarly-sized gastropods (Novak, 2013)), and structural defenses (e.g., the pulmonate limpet *Siphonaria australis* with its mucous-rich foot is typically drilled while patellid limpets like *Cellana ornata* are simply flipped (Fig. 1, *pers. obs.*)). For such fundamental aspects of biology to dramatically change in a way that reduces variation over ecological time-scales seems unlikely (but see Thompson, 1998; and many others).

The greatest weakness of the above-argued reason for *H. haustorium*’s feeding-rate consistency is therefore my inference that its detection times remained positively correlated between time periods (i.e. *r_w,z_ >* 0 in eqn. 4), this being not only a matter of the species’ biological attributes but also of *H. haustorium*’s behavioural prey choices and predatory tactics, which are likely to be far more labile^2^ (Blomberg *et al*., 2003). More specifically, although I did not assume a given species’ detection time was the same between time periods, I did assume that whelks of a given size would exhibit the same detection time for a prey of a given identity and size at a given temperature. I thereby allowed for each of these variables to differ from observation to observation, site to site, and across time periods, assuming only their relationship to detection times to have remained unchanged. This assumption seems defensible given the relatively slow-to-evolve physiological and structural basis of whelk handling times (Carriker, 1981): rasping and digesting and involving the evolutionary arms race between whelks and their prey. However, handling and hence detection times may be far more changeable for other types of predator-prey interactions depending on the species’ biological attributes and aspects of the feeding process on which feeding surveys rely (e.g., whether feeding events are observed directly or by the examination of gut contents (Novak *et al*., 2017)). For some species, such as those involving more specialized predator-prey pairs (DeLong & Coblentz, 2021), handling times could be just as labile as species abundances and prey preferences, and could in fact respond to these as well (Okuyama, 2010; Stouffer & Novak, 2021). In such contexts where the consistency of detection times may be weak, detection-time variation will need to be large for the statistical mechanism of correlated ratios to contribute to feeding-rate stability.

Two additional considerations pertain more to methodological details. First, it *is* possible for a spurious correlation to occur when evaluating feeding-rate stability through diet surveys. This is because the apparent diet proportions (*n_i_/n* of eqn. 1) will themselves become correlated if the sample sizes (*n*) of both sets of surveys are correlated, just as Pearson (1897) warned. This was not the case in this study (Table 1; *r* = 0.01, *p* = 0.98; *r*_10_ = *−*0.28, *p* = 0.65; *r_s_* = *−*0.40, *p* = 0.52), but may be quite likely to occur in other studies when sites exhibit a consistent gradient in predator abundances due to underlying environmental or productivity differences (e.g., Novak, 2013; Winemiller, 1990). Second, although it is possible that the higher overall feeding rate of *H. haustorium* in 1968–9 versus 2004 was due to a change in their size distribution, it is also possible that Paine’s and my feeding surveys differed in a biased way in regards to our ability to find larger versus smaller, or feeding versus non-feeding individuals; on average, Paine was almost 2.5 times more likely to find feeding individuals than me (Table 1). Given Paine’s extensive experience with intertidal feeding surveys, the fact that he and his frequent field assistant, Terrence Beckett, compared and saw no difference between their independent surveys^3^, and the fact that smaller and non-feeding individuals tend to be more difficult to locate (especially by relative novices like me in 2004), I consider biases due to differences in survey ability improbable. The issue of bias in resurvey studies more generally requires attention nonetheless, just as it does when manipulative experiments are repeated (Kimmel *et al*., 2021).

## Conclusions

Overall, the results of my study draw attention to the potential for the detection times of feeding events to alter the interpretation of predator diet data. Variation in detection times has been little studied relative to the substantial effort that has gone into the study of foraging strategies and prey preferences. Most relevant work has focused on the gut-evacuation rates of prey mass in fishes, but with little focus on generalist predators’ diverse prey attributes per se (Preston *et al*., 2017). In the functional-response literature, handling and digestion times are primarily considered to be important only at high prey abundances where feeding rates are limited by saturation or satiation (Jeschke *et al*., 2002). The potential for the effect of which Pearson (1897) warned to alter the interpretation of apparent diets for many more types of taxa indicates that more attention to detection times is warranted, and that factors to which handling and digestion times are sensitive may be more important in structuring feeding rates (i.e. process-rate variation) than currently assumed, even at low prey abundances. Feeding rates may be far less changing than inferred by surveys of apparent diets and community structure alone. As such, an improved understanding of detection times will not only be relevant to historical resurveys and other temporal analyses of community and interaction-network structure (Bramon Mora *et al*., 2020), but will also be relevant to studies describing biogeographic patterns in these to infer how communities function (Bartley *et al*., 2019; Tylianakis & Morris, 2017).

## Acknowledgments

I am grateful to Bob Paine (1933–2016) for providing his data to me before the first field season of my Ph.D. work in 2004. I also thank William (Bill) J. Ballantine (1937–2015) for his hospitality while I was at the Leigh Marine Laboratory. As recorded in his notebooks, Paine received substantial field assistance from Terrence (Terry) Beckett, as well as from Dick Martin, Charles (Chuck) Galt, M. Larkum, and Dreseks (first names unknown). My field season was enabled by the support of the University of Chicago Hinds Fund. The subsequent laboratory-based detection time experiments were enabled by a NSF DEB-0608178 award and an EPA STAR fellowship. The approach to feeding rate estimation was derived with the support of NSF DEB-1353827. This manuscript was completed while on sabbatical as a visiting scholar hosted by Jennifer Ruesink at the University of Washington. I thank Kyle Coblentz, Bruce Menge, and Rebecca Terry for their valuable comments on the manuscript.

## Code and data availability

All code and data used in the presented analyses, as well as data which Bob collected at additional sites to which I was unable to return, are available at https://github.com/marknovak/ NZPaineFrates.

## Footnotes

1 Paine’s notebook records his having measured the shells of 15 large individuals, 65.0, 65.5, 65.7, 66.2, 67.5, 68.0, 68.3, 68.5, 68.6, 69.4, 69.6, 71.9, 73.1, 74.0 and 76.8 mm in length, in a Maori midden of unknown age found somewhere between North Cape (Otou) and Parengarenga Harbour.

2 Anecdotally, populations of *H. haustorium* around Kaikoura on the east coast of the South Island, where mussels are rare, could not be brought to feed on them in the lab (although rare field observations thereof occurred), while populations from the west coast, where mussels are abundant, readily did so (Novak, 2008).

3 As recorded in Paine’s field notes.

## Notes

https://github.com/marknovak/NZPaineFrates

### Competing Interest Statement

The authors have declared no competing interest.

