## Supplementary Materials for "High variation in handling times confers 35-year stability to predator feeding rates despite altered prey abundances and apparent diet proportions"

Mark Novak

### Contents

**S1 Supplementary tables** **2**

**S2 Regression summaries** **4**

**S3 Additional measures of diet and community similarity** **5**

**S4 Spurious versus non-spurious correlations of ratios** **7**

**References** **9**

### S1 Supplementary tables

Table S1: The locations where Paine and I surveyed *Haustrum haustorium*'s diet and the abundances of its prey in 1968-9 and 2004 for which data are posted to the public repositories indicated in the main text. Missing coordinates are unknown.

| | | | **Feeding** | | **Abundance** | |
|---|---|---|---|---|---|---|
| Site | Latitude | Longitude | 1968-9 | 2004 | 1968-9 | 2004 |
| Waikuku Bay | −34.4720 | 173.0079 | x | | | |
| Leigh - Waterfall Rocks | −36.2688 | 174.8060 | x | x | x | x |
| Leigh Goat Island Reserve | −36.2688 | 174.8060 | x | | | |
| Leigh - Echinoderm Reef | −36.2696 | 174.7937 | x | x | | |
| Leigh - Tabletop Rocks and Boulders | −36.2701 | 174.8025 | x | x | | |
| Leigh Harbour | −36.2881 | 174.8080 | x | | | |
| Red Beach - Whangaparaoa | −36.6007 | 174.7092 | x | x | x | x |
| Rangitoto Island - Whites Beach | −36.7754 | 174.8334 | x | x | x | x |
| Takapuna | −36.8160 | 174.8087 | x | | | |
| Kaikoura Paine's | | | x | | | |
| Waikukua Bay-2 | | | x | | | |
| Whangarei | | | x | | | |
| Tapotupotu Bay West | −34.4347 | 172.7129 | | x | | |
| Leigh Shadow Rocks | −36.2715 | 174.8091 | | x | | |
| Tungutu Point | −36.5075 | 174.7231 | | x | | |
| Red Beach Cliff | −36.6003 | 174.7075 | | x | | |
| Opunake | −39.4597 | 173.8485 | | x | | |
| Pourere Tuingara Point | −40.1376 | 176.8650 | | x | | |
| Castle Point Cave | −40.8997 | 176.2310 | | x | | |
| Castle Point Boulders | −40.9006 | 176.2302 | | x | | |
| Island Bay Lab Rocks | −41.3490 | 174.7649 | | x | | |
| Matakitakiakupe Cape Palliser | −41.6125 | 175.2742 | | x | | |
| Cape Foulwind NWPlatform | −41.7461 | 171.4666 | | x | | |
| Cape Foulwind | −41.7526 | 171.4586 | | x | | |
| Tauranga Bay North | −41.7653 | 171.4560 | | x | | |
| Tauranga Head | −41.7738 | 171.4555 | | x | | |
| Tauranga Head West | −41.7764 | 171.4514 | | x | | |
| Tauranga Head SWcorner | −41.7768 | 171.4523 | | x | | |
| Ward Beach | −41.8483 | 174.1836 | | x | | |
| Charleston Joyce Bay | −41.9022 | 171.4350 | | x | | |
| Memorial Garden Rocks | −42.4044 | 173.6851 | | x | | |
| Whakatu Point | −42.4143 | 173.7062 | | x | | |
| Avoca Point North | −42.4161 | 173.7076 | | x | | |
| Lighthouse Reef | −42.4239 | 173.7169 | | x | | |
| First Bay | −42.4261 | 173.7143 | | x | | |
| Limestone Bay Point | −42.4267 | 173.6872 | | x | | |
| Raramai | −42.4586 | 173.5520 | | x | | |
| Oaro South | −42.5239 | 173.5050 | | x | | |

Table S2: Prey for which *Haustrum haustorium*'s prey-specific detection times had not been measured in the laboratory experiments of Novak (2013) were assigned the regression coefficients of prey species for which they had been measured.

| **Unmeasured** | | **Matched to measured** | |
|---|---|---|---|
| Predator | Prey | Predator | Prey |
| H. haustorium | Atalacmea fragilis | H. haustorium | Cellana radians |
| H. haustorium | Cellana stellifera | H. haustorium | Cellana radians |
| H. haustorium | Chamaesipho columna | H. haustorium | Chamaesipho spp |
| H. haustorium | Cominella adspersa | H. haustorium | Haustrum scobina |
| H. haustorium | Crassostrea gigas | H. scobina | Mytilus galloprovincialis |
| H. haustorium | Dicathais orbita | H. haustorium | Haustrum scobina |
| H. haustorium | Diloma bicanaliculata | H. haustorium | Diloma aethiops |
| H. haustorium | Diloma nigerrima | H. haustorium | Diloma aethiops |
| H. haustorium | Diloma zelandica | H. haustorium | Diloma aethiops |
| H. haustorium | Fossarina rimata | H. haustorium | Risellopsis varia |
| H. haustorium | Haustrum haustorium | H. haustorium | Haustrum scobina |
| H. haustorium | Mytilus galloprovincialis | H. scobina | Mytilus galloprovincialis |
| H. haustorium | Nerita atramentosa | H. haustorium | Diloma aethiops |
| H. haustorium | Notoacmea parviconoidea | H. haustorium | Notoacmea spp |
| H. haustorium | Paratrophon patens | H. haustorium | Haustrum scobina |
| H. haustorium | Trimusculus conicus | H. haustorium | Siphonaria australis |
| H. haustorium | UNID Chiton | H. haustorium | Plaxiphora caelata |
| H. haustorium | UNID Diloma | H. haustorium | Diloma aethiops |
| H. haustorium | UNID Limpet | H. haustorium | Notoacmea spp |
| H. haustorium | UNID Notoacmea | H. haustorium | Notoacmea spp |
| H. haustorium | UNID Snail | H. haustorium | Diloma aethiops |
| H. haustorium | Zeacumantus subcarinatus | H. haustorium | Austrolittorina cincta |

### S2 Regression summaries

Table S3: Summary table for the regression of predator size on prey size.

| | Estimate | Std. Error | t value | Pr(>—t—) |
|---|---|---|---|---|
| (Intercept) | 2.336 | 0.0466 | 50.1 | $9.17e-170$ |
| log(PreySize) | 0.456 | 0.0182 | 25.1 | $9.61e-83$ |

Table S4: Summary table for the regression of predator size on prey size and time period (*Year*).

| | Estimate | Std. Error | t value | Pr(>—t—) |
|---|---|---|---|---|
| (Intercept) | 2.670792 | 1.307244 | 2.043 | $4.17e-02$ |
| log(PreySize) | 0.453821 | 0.020303 | 22.352 | $3.00e-71$ |
| Year | −0.000166 | 0.000647 | −0.257 | $7.98e-01$ |

Table S5: Summary table for the regression of predator size on prey size, time period (*Year*), and their interaction.

| | Estimate | Std. Error | t value | Pr(>—t—) |
|---|---|---|---|---|
| (Intercept) | 0.491060 | 5.83524 | 0.0842 | 0.933 |
| log(PreySize) | 1.357255 | 2.35701 | 0.5758 | 0.565 |
| Year | 0.000932 | 0.00294 | 0.3174 | 0.751 |
| log(PreySize):Year | −0.000456 | 0.00119 | −0.3833 | 0.702 |

Table S6: Summary table for the regression of prey-specific feeding rate on prey-specific abundance, time period (*Year*), and their interaction.

| | Estimate | Std. Error | t value | Pr(>—t—) |
|---|---|---|---|---|
| (Intercept) | 15.36190 | 30.2353 | 0.508 | 0.617 |
| log10(N.mean) | −9.84166 | 21.3753 | −0.460 | 0.650 |
| Year | −0.00888 | 0.0152 | −0.583 | 0.566 |
| log10(N.mean):Year | 0.00522 | 0.0108 | 0.485 | 0.633 |

Table S7: Summary table for the regression of prey-specific feeding rate on prey-specific abundance and time period (*Year*).

| | Estimate | Std. Error | t value | Pr(>—t—) |
|---|---|---|---|---|
| (Intercept) | 2.18267 | 13.03154 | 0.167 | 0.86851 |
| log10(N.mean) | 0.52616 | 0.18387 | 2.862 | 0.00907 |
| Year | −0.00224 | 0.00657 | −0.341 | 0.73652 |

Table S8: Summary table for the regression of prey-specific feeding rate on prey-specific abundance.

| | Estimate | Std. Error | t value | Pr(>—t—) |
|---|---|---|---|---|
| (Intercept) | $-2.257$ | 0.254 | $-8.89$ | $6.70e-09$ |
| log10(N.mean) | 0.521 | 0.180 | 2.90 | $8.08e-03$ |

### S3 Additional measures of diet and community similarity

The correlation and distance-based comparisons of the main text included only prey species which both Paine and I observed *H. haustorium* feeding on at a given site. To compare *H. haustorium*'s apparent diet and each site's community structure between time periods more generally (i.e. including species incidences), I performed additional analyses that also (*i*) included prey species which only one of us observed in our feeding surveys and (*ii*) included prey species which only one of us observed as well as additional (non-prey) mobile species which we observed in our abundance surveys.

To assess time-period similarities in diet and prey abundances at all five sites where both Paine and I performed feeding surveys, I used the classic incidence-based Jaccard index ($J_{class}$), the abundance-based Jaccard index ($J_{abd}$), and the estimator for the abundance-based Jaccard index ($\hat{J}_{abd}$) (Chao *et al.*, 2005). While $J_{class}$ quantifies compositional similarity (species overlap), $J_{abd}$ reflects the probability that two randomly chosen individuals, one from each time period, both belong to any of the shared species seen in both time periods (not necessarily to the same shared species). The estimator $\hat{J}_{abd}$ attempts to account for shared but rare species that were not observed due to incomplete sampling. Overall, these analyses indicate low to intermediate levels of similarity in the composition of *H. haustorium*'s apparent diet and community that were driven by changes in the occurrence of low-frequency prey/species; for most sites, similarities were higher when considering prey frequency/species abundance and differed little between $J_{abd}$ and $\hat{J}_{abd}$ (Table S9).

To visualize similarities in community structure, I performed a two-dimensional non-metric multi-dimensional scaling analysis with the *vegan* R-package (Oksanen *et al.*, 2020) using the Bray-Curtis metric to quantify distances between surveyed quadrats. This analysis indicated that all three sites surveyed by both Paine and me have changed in their community structure, with my surveys indicating more similar communities (both within and between sites) than did Paine's surveys (Fig. S1).

Table S9: The between time period similarity of *Haustrum haustorium*'s apparent diet – at the sites where Paine and I performed either feeding surveys only or both feeding and abundance surveys – as quantified by the incidence-based Jaccard index ($J_{class}$), as well as the abundance-based Jaccard index ($J_{abd}$) and the estimator for the abundace-based Jaccard index ($\hat{J}_{abd}$).

| | **Feeding observations** | | | **Prey abundances** | | |
|---|---|---|---|---|---|---|
| Site | $J_{class}$ | $J_{abd}$ | $\hat{J}_{abd}$ | $J_{class}$ | $J_{abd}$ | $\hat{J}_{abd}$ |
| Leigh - Echinoderm Reef | 0.50 | 0.78 | 0.80 | - | - | - |
| Leigh - Tabletop Rocks and Boulders | 0.38 | 0.30 | 0.32 | - | - | - |
| Leigh - Waterfall Rocks | 0.50 | 0.87 | 0.98 | 0.77 | 0.92 | 0.92 |
| Rangitoto Island - Whites Beach | 0.50 | 0.74 | 0.81 | 0.33 | 0.45 | 0.45 |
| Red Beach - Whangaparaoa | 0.25 | 0.74 | 0.74 | 0.4 | 0.81 | 0.81 |

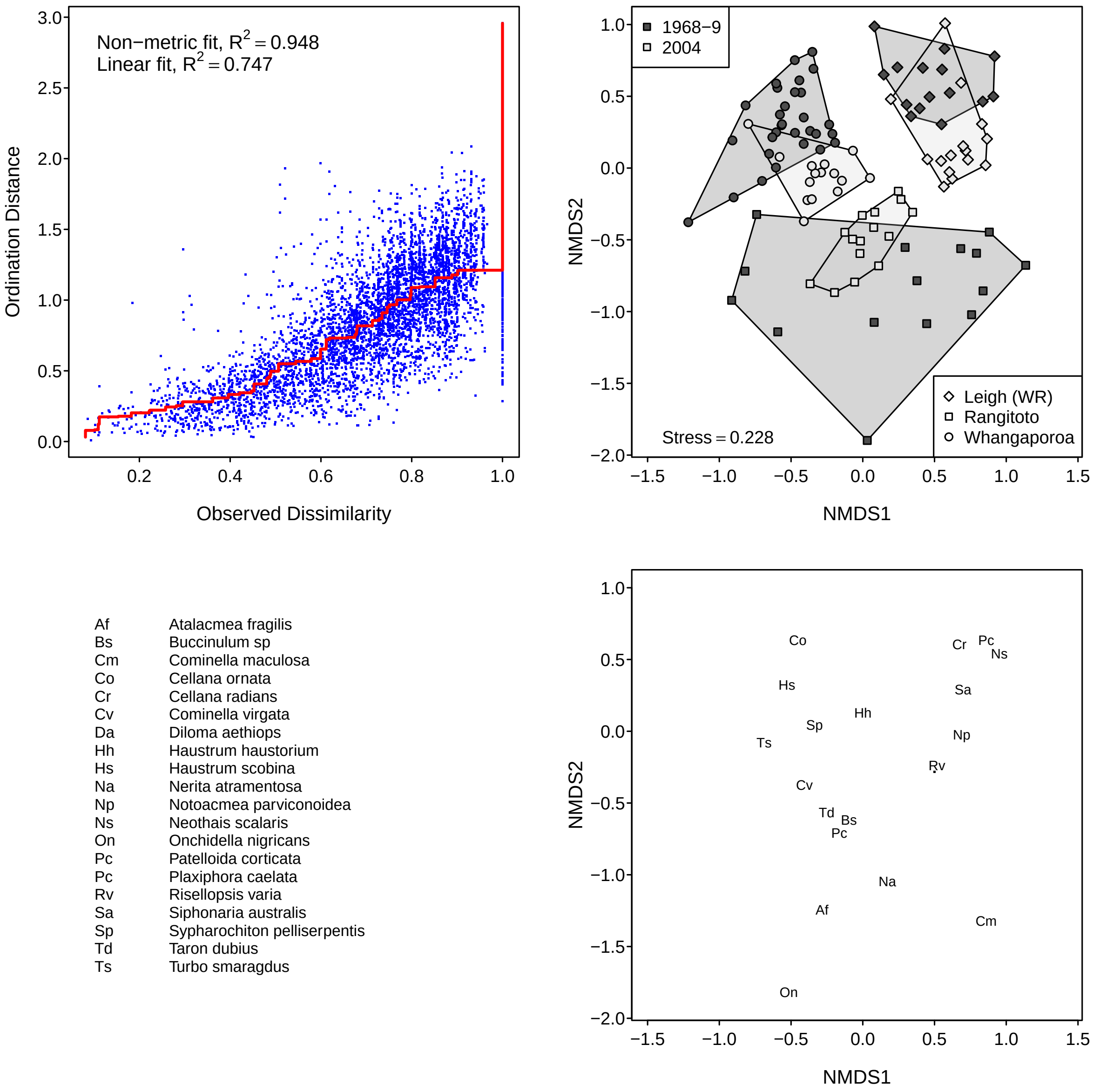


Figure S1: Non-metric multi-dimensional scaling using Bray-Curtis distances between quadrat-specific mobile species counts (including *H. haustorium*'s prey and other, non-prey species) at the three sites which Paine surveyed for species abundances in 1968–9 and which I resurveyed in 2004.

### S4 Spurious versus non-spurious correlations of ratios

The difference between spurious and non-spurious interpretations of the correlation of ratios may be illustrated using variable permutations, as shown by the following `R` script (also available at `https://github.com/marknovak/NZPaineFrates/blob/main/code/RatioCorr.R`).

```
###################################################
###################################################
# Spurious versus non-spurious ratio correlations
###################################################
###################################################
library(MASS) # for mvrnorm
#~~~~~~~~~~~~~~~~~~~~~~~~~~~~~~~~~~~~~~~~
# Multivariate-normal random variables
# independent numerators
# but correlated denominators
#~~~~~~~~~~~~~~~~~~~~~~~~~~~~~~~~~~~~~~~~
S <- 100 # Sample size
xn <- rnorm(S, 10, 0.5) # first numerator
yn <- rnorm(S, 10, 0.5) # second numerator
cov <- 0.9 # Covariance between denominator variables
d <- data.frame(mvrnorm(n = S,
mu = c(xd = 10, yd = 10),
Sigma = rbind(c(1, cov), c(cov, 1))))
xd <- d$xd # first denominator
yd <- d$yd # second denominator

# Only the denominators are correlated
pairs(cbind(xn, yn, xd, yd))
cor(cbind(xn, yn, xd, yd))

# The ratios are correlated
cor.obs <- cor(xn/xd, yn/yd)

# Contrast spurious versus non-spurious inferences be permuting either
# just the numerators or both the numerators and denominators

num.sim <- 9999 # number of permutations

# It's a spurious correlation when drawing inference about the numerators
cor.spur <- replicate(num.sim, cor(sample(xn) / xd,
(yn) / yd))
# But *not* a spurious correlation when drawing inference about the ratios
cor.nonspur <- replicate(num.sim, cor(sample(xn/xd),
yn/yd))

cor.obs
```

```
mean(cor.spur)
mean(cor.nonspur)

# Inspect distributions of observed correlations
h1 <- hist(cor.nonspur, breaks = 200, xlim = c(-1,1),
main = '', xlab = 'Correlation')
abline(v = cor.obs, col = 'blue', lwd = 2)
abline(v = mean(cor.spur), col = 'red', lwd = 2)
h2 <- hist(cor.spur, breaks = 200, xlim = c(-1,1),
add = TRUE)
legend('topleft',
legend = c('Observed',"Mean (Expected)"),
lty = 1,
col = c('blue','red'),
bty = 'n')

# Conclusion: In the context of comparing feeding rates,
#  we are drawing inference about the correlation of the ratios
#  (not the numerator diet proportions).
# A correlation of zero,  mean(cor.nonspur),
#  is thus the appropriate null hypothesis.
# If we were drawing inference on the numerator diet proportions,
#  then the non-zero correlation, mean(cor.spur),
#  is the appropriate null hypothesis.
```

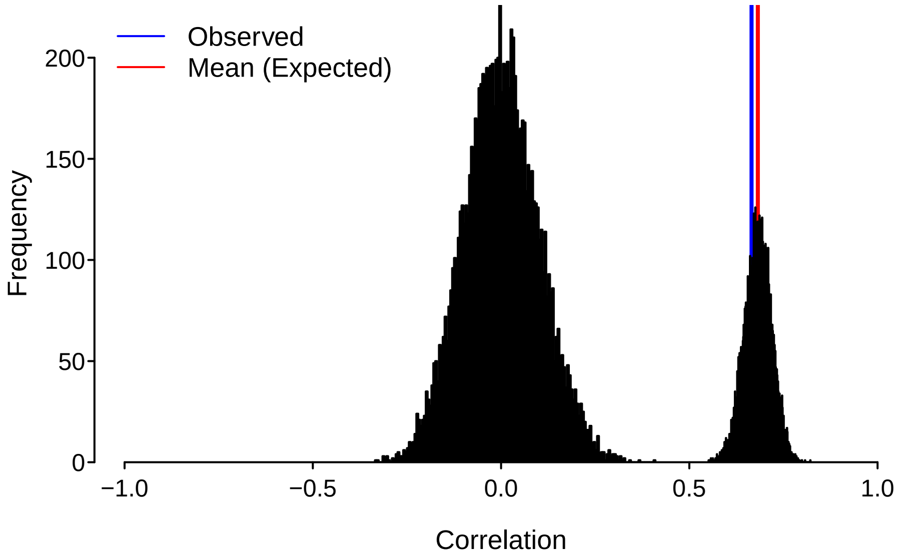


Figure S2: Frequency histograms for the correlation values when the numerators are permuted but the denominator variables remain correlated (right distribution) versus when the ratios themselves are permuted (left distribution). (Produced by above `R` code.)
